# Haplotypes associated to gene expression in breast cancer: can they lead us to the susceptibility markers?

**DOI:** 10.1101/248047

**Authors:** Hege Edvardsen, Bettina Kulle, Anya Tsalenko, Grethe Irene Grenaker Alnӕs, Fredrik Ekeberg Johansen, Espen Enerly, Aditya Vailaya, Per-Eystein Lønning, Åslaug Helland, Ann-Christine Syvänen, Zohar Yakhini, Anne-Lise Børresen-Dale, Arnoldo Frigessi, Vessela N. Kristensen

## Abstract

We have undertaken a systematic haplotype analysis of the positional type of biclusters analysing samples collected from 164 breast cancer patients and 86 women with no known history of breast cancer. We present here the haplotypes and LD patterns in more than 80 genes distributed across all chromosomes and how they differ between cases and controls. We aim by this to 1) identify genes with different haplotype distribution or LD patterns between breast cancer patients and controls and 2) to evaluate the intratumoral mRNA expression patterns in breast cancer associated particularly to the cancer susceptibility haplotypes. A significant difference in haplotype distribution between cases and controls was observed for a total of 35 genes including *ABCC1, AKT2, NFKB1, TGFBR2* and *XRCC4*. In addition we see a negative correlation between LD patterns in cases and controls for neighboring markers in 8 genes such as *CDKN1A, EPHX1* and *XRCC1*.

## Introduction

The common disease common variant hypothesis is the foundation for large scale whole genome analyses of extensive population cohorts aiming at identifying low penetrant markers that in concert result in an increased risk of cancer. Nearly 1300 published GWAS studies have so far identified 6551 markers associated with various diseases and traits such as asthma, multiple sclerosis and various cancer types (1). For breast cancer specifically SNPs in 41 genes including ***FGFR2, TOX3, TERT*** and ***ERBB4*** have been associated to the disease (***2-5***). These studies view the risk of common genetic variation only and the number of markers is restricted to the number of SNPs on the studied arrays without focus on particular genes or functionality. Here we have taken an alternative route based on a candidate gene approach without restriction to frequency. Moreover, we did not study single disease associated SNPs but looked for differences in haplotype distribution and LD patterns between cases and controls. Linkage disequilibrium (LD) is the association between two markers (SNPs) resulting from common inheritance of two typically nearby loci. LD is eroded by mutations, gene conversions and recombination events, and is influenced by the age of the mutations as well as the history and size of the populations in which they are studied. Several measurements are used to estimate LD such as D’ (6) and r^2^ (7). D’ shows larger variability within and between populations and is more influenced by sample size (8,9). D’ takes into account the history of the markers and is more robust with regards to frequency while r^2^ is less affected by problems related to sampling (8). It is also possible to use statistical estimates of population recombination rates (ρ) instead of pairwise measures of LD (9). This measure correlates well across populations and relates the LD pattern directly to the underlying recombination process (7). Haplotypes are strings or combinations of co-inherited SNPs residing at regions of high LD and separated by areas with high recombination and low LD (8). They are inherited from parents as a single unit and tend to break at recombination hotspots (3). In population studies in contrast to linkage analysis in families, an absolute determination of haplotypes is not possible, but studies of phased estimations have proven these to be a very good approximation. The results from these studies indicate an error in assigning phase to genotypes of approximately 5 % in unrelated individuals (10). This uncertainty can be adjusted for as we have previously described (11).

The choice of LD block was motivated by our studies in eQTLs. Our findings indicate that the breast cancer risk variants found by the GWASs may exert their effect through the regulation of expression, and that the genes harboring these risk variants are significantly differentially expressed between the well established breast cancer subtypes (12). Given the significant role mRNA expression patterns play in the development of breast cancer, we hypothesize that SNPs associated to clusters of deregulated co-expressed mRNA transcripts may lead us to novel susceptibility markers. We have previously described that among 583 candidate SNPs in 203 genes of the reactive oxygen species metabolism/signaling, there are SNPs significantly associated over the random to the expression of subsets of unselected transcripts in the tumor of breast cancer (13). Furthermore, these subsets of transcripts are enriched for given functional pathways also over the random. Multiple SNPs (biclustes) that together share significantly many common associations to a set of transcripts were identified. These biclusters were either located in different genes on different chromosomes, suggesting a multi-locus regulatory effect on a pathway (functional biclusters) or clustered in the same gene or chromosomal region (positional biclusters). With the present study we have undertaken a systematic haplotype analysis of these positional biclusters extending the analysis to samples collected from 164 breast cancer patients and 86 women with no known history of breast cancer. We aim by this study to 1) formally assess the degree of LD between the SNPs in the positional biclusters associated to expression 2) use these eQTL hits to identify cancer susceptibility haplotypes by comparing the distribution or LD patterns between 1592 breast cancer patients and 1892 controls.

## Material and Methods

### Genotyping

We have genotyped 164 breast cancer patients and 86 healthy women with no known history of cancer (two negative mammography screenings). The panel of SNPs genotyped are thoroughly described in (14) but in short, SNPs in candidate genes involved in the metabolism of reactive oxygen species and xenobiotics, DNA repair, cell cycle and apoptosis were genotyped using a mini-sequencing (SNP-IT) method multiplexing up to twelve SNPs in one tube. The polymerase chain reaction, clean up with ExoI and SAP and SNP-IT reaction are performed in one tube and the reaction mix hybridized to an array. Each of the twelve SNP-IT primers contain a tag that utilizes sorting of the multiplex reaction on the array. The mini-sequencing reaction is a two colour reaction and signal is detected after laser excitation of the fluorophores on the SNP-stream UHT system.

### Validation analysis of selected SNPs

Validation of selected SNPs were done using the Sequenom MassARRY platform and iPLEX genotyping assays (http://www.sequenom.com/home/) (15).

### Microarray expression analysis

For 50 of the breast cancer patients, expression data were also available. Tumour tissue (20-50mg) was dissected and powdered in liquid nitrogen and total RNA was prepared by standard procedures. Whole genome microarray expression analysis has been performed using cDNA microarrays as described in (16-18).

### SNP-expression association analysis

Unselected subset of 3351 mRNA transcripts was obtained by filtering for signal quality (ratio of spot intensity over background exceeding 1.5 in at least 80% of the experiments in each dye channel). The analysis of the SNP-expression associations are published earlier in (13). For these patients additional 28 SNPs were available for the haplotype analysis. In short, the correlation between genotypes and expression level of the different mRNA transcripts were assessed using three, different statistical approaches; ANOVA, QMIS and LOOCV. For each SNP locus and each transcript, the one-way ANOVA p-value was computed for the expression vector and grouping of the samples based on SNP locus genotypes (19) assuming the null hypothesis that the expression level distributions are the same, regardless of the genotype class. QMIS (Quantitative Mutual Information Score). For a SNP locus s and an expression vector q of transcript t, let G be a partition of samples induced by the genotype values at locus s. For an expression level threshold p, let Cp be a partition of samples defined by the q<p and q≥p. The mutual information score (MIS) is the difference between the entropy of the partition Cp and the conditional entropy of Cp given G: MIS(Cp, G) = H(Cp) - H(Cp |G), where H is the entropy function. The quantitative mutual information score is defined to be the maximum possible MIS, i.e., QMIS(C,G) = maxmin(q) ≤p≤max(q)MIS(Cp, G). An exact p-value for the mutual information score can be computed exactly by an efficient exhaustive approach (20). In this case, the null hypothesis is that genotype values have the same distribution, regardless of expression levels. For QMIS, 769 SNP-transcript association pairs with p-values ≤ 1.0E -04 were observed, representing an FDR of 0.2. LOOCV (Leave Out Cross Validation) for a given SNP in the data set, its genotypes were utilized to group samples. For each grouping, leave-one-out-cross-validation analysis was performed, trying to predict from the expression data which genotype group each sample belongs to (similar to the methods described in (21)).

### Gene Ontology analysis (GO)

The group of transcripts associated to the same SNP or group of SNPs was analysed with regards to enrichment of GO terms based on GO terms downloaded from Source (http://source.stanford.edu/cgi-bin/source/sourceSearch), for this analysis the p-value cut-off for the SNP-expression association was set at 0.05 and 0.01. The significant overrepresentation for a GO term was calculated taking into account the total number of; 1) genes on the expression array, 2) genes associated with the GO term, 3) genes associated to the SNP and 4) the number of genes associated with the SNP or group of SNPs, that belong to the GO term. The z score was calculated according (14) by subtracting the expected number of genes in a GO term from the observed and diving this by the standard deviation of the observed.

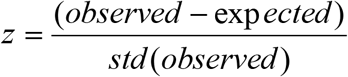

### Calculation of LD and Spearman’s correlation coefficient

SNPs that had discovery rate lower than 75% were excluded. Initially, the panels of SNPs were screened for clusters containing a minimum of 3 SNPs with no more than 100 kb between neighboring SNPs. For the genes represented in these clusters - all genotyped SNPs were included. LD estimations were done in two steps. First, we estimated the haplotypes for cases and controls separately from our population genotype data using the recombination model implemented in the program PHASE (Stephens, M. et al.) with 5 different seeds and 100. The significance of the difference in haplotype distribution between cases and controls was calculated in Phase. The second step was the evaluation of the LD for all included genes. For this purpose, we calculated pairwise D’ for cases and controls separately for all possible SNP combinations within a gene and under consideration of the uncertainty in phase estimation (11). PHASE also provides the recombination rate as a measure of dependency between the SNPs for all adjacent SNPs within a gene. To evaluate the difference for each gene between the LD-patterns of cases and controls, we calculated Spearman’s correlation coefficient ρ as done in (9). The correlation is given as a value between -1 ≤ 0 ≤1, where 0 indicates no correlation, whereas -1 and 1 indicates high negative and positive correlation respectively. We calculated this nonparametric correlation coefficient 1) using all markers for D’ and 2) using only adjacent markers for ρ.

### Analysing the relationship between haplotypes and expression levels of transcripts associated to multiple SNPs within a gene

The non-parametric Mann Whitney or Kruskal Wallis test was used to analyse the possible connection between the haplotypes estimated for a gene and the expression levels of transcripts associated to all or a subset of the SNPs within the given haplotypes. The analysis were performed using SPSS v15.0, the p-values are exact (50 iterations), two-tailed and not corrected for multiple testing.

### Estimating population subdivision - calculating the fixation index

Population subdivision was estimated using the Arlequin Software to calculate the Fixation index (Fst). This index measures the population differentiation between two groups and its values range from 0 to 1 (with 0 meaning that the populations are completely similar with regard to allele frequencies and 1 being that the populations are completely differentiated (22).

## Results and discussion

The overall study design is given in **Supplementary Figure 1.** A total of 687 SNPs in 203 genes selected from pathways related to the ROS metabolism and signaling were genotyped in 169 breast cancer patients and 86 controls (*14).* Haplotypes were inferred and of the 687 SNPs, a subset of 457 SNPs were available at HapMap with associated frequency information. The full list of SNPs used in the analysis can be found in **Supplementary Table 1** together with information on gene affiliation, chromosomal position, allelic variation and strand genotyped.

### Impact of multiple SNPs (biclusters) on the expression profile;

For 50 of the patients genotyped here expression data were available and we have previously reported the association of 538 SNPs to the intratumoral mRNA expression in these patients (13). Many of the studied genes, e.g. *ABCC1, ALOX12, DPYD, GSTM3, NOX3, IL10* and *IL8* were shown to harbor multiple SNPs significantly associated to the level of transcripts *in cis* and *trans* (for full list see **Supplementary Table 2).** We have formally assessed the degree of LD between the multiple SNPs regulating the same group of transcripts and observe that many of these are in strong linkage disequilibrium such as in the genes of *DPYD, TXNIP, GSTA4, PPP1R9A, NFKBIA, IGF1R, ABCC1* and as shown for *XDH* and *IL1R1* on chromosome 2 **Figure 1** (figures for all other chromosomes are given in **Supplementary Figure 2a-u).** Further analyzing the characteristics of these subsets of coexpressed transcripts by gene ontology analysis (p-value cut-off for the SNP-transcript association: 0.01), we find for SNPs in more than 25 genes a significant overrepresentation of GO terms in the list of regulated transcripts at p-value< 0.001 **(Table 1).** Compelling examples are: 1) 18 SNPs in DPYD (involved in pyrimidine base degradation) which together with a SNP in *GSTM4* all are associated to the expression of a group of 10 transcripts among which there is an overrepresentation of the GO term *regulation of cell growth* and 2) 6 SNPs in *GSTA4* associated to a group of 20 transcripts with an overrepresentation of the GO term *transcriptional activator activity.*

**Figure 1.**
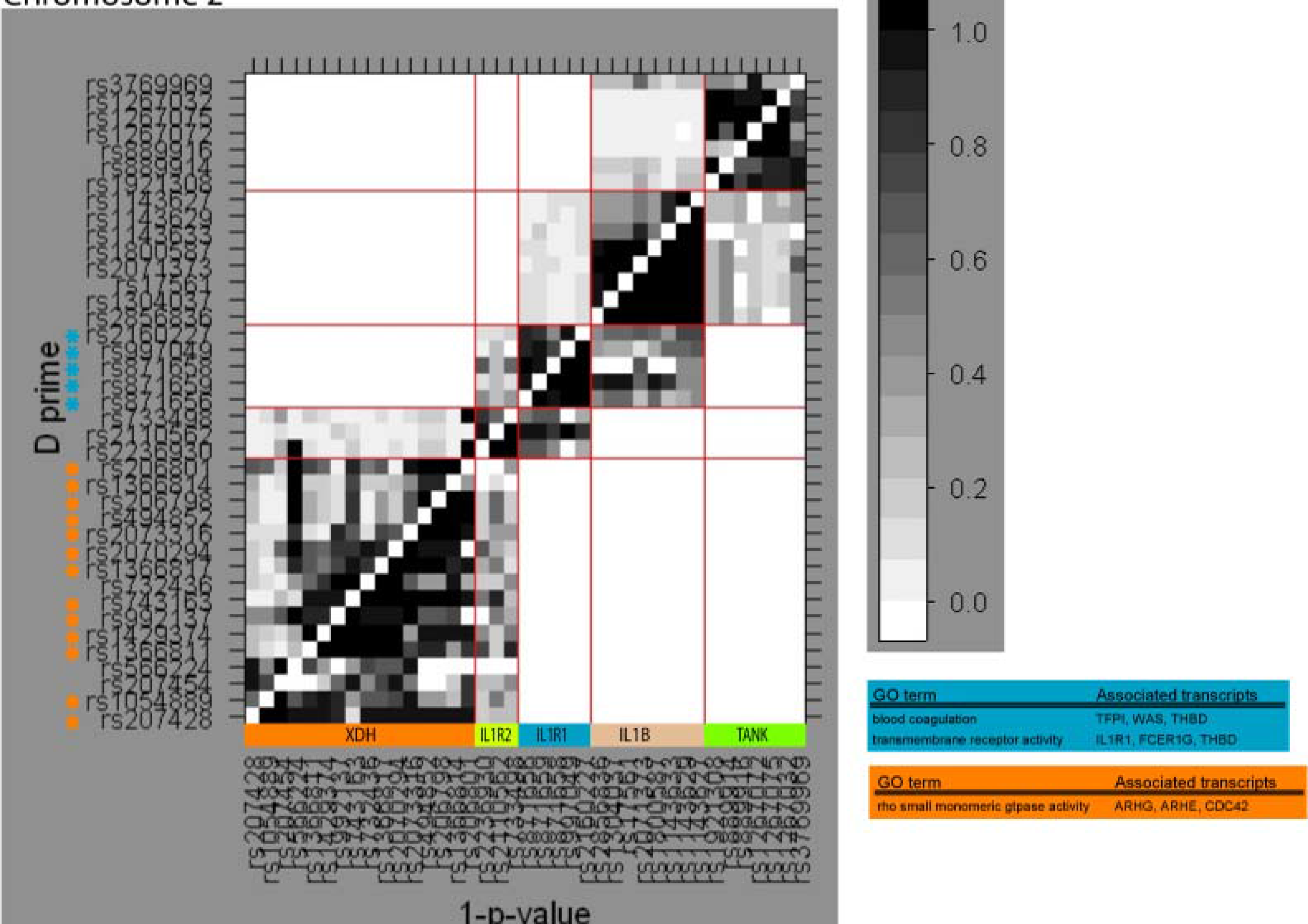
LD pattern in chromosome 2 for the cases together with information on overrepresented GO terms among associated transcripts. X-axis indicates the significance level of the LD while the |D’| values are plotted on the Y-axis, values along the diagonal are intragenic, adjacent panels give information on intergenic regions.

**Table 1.**
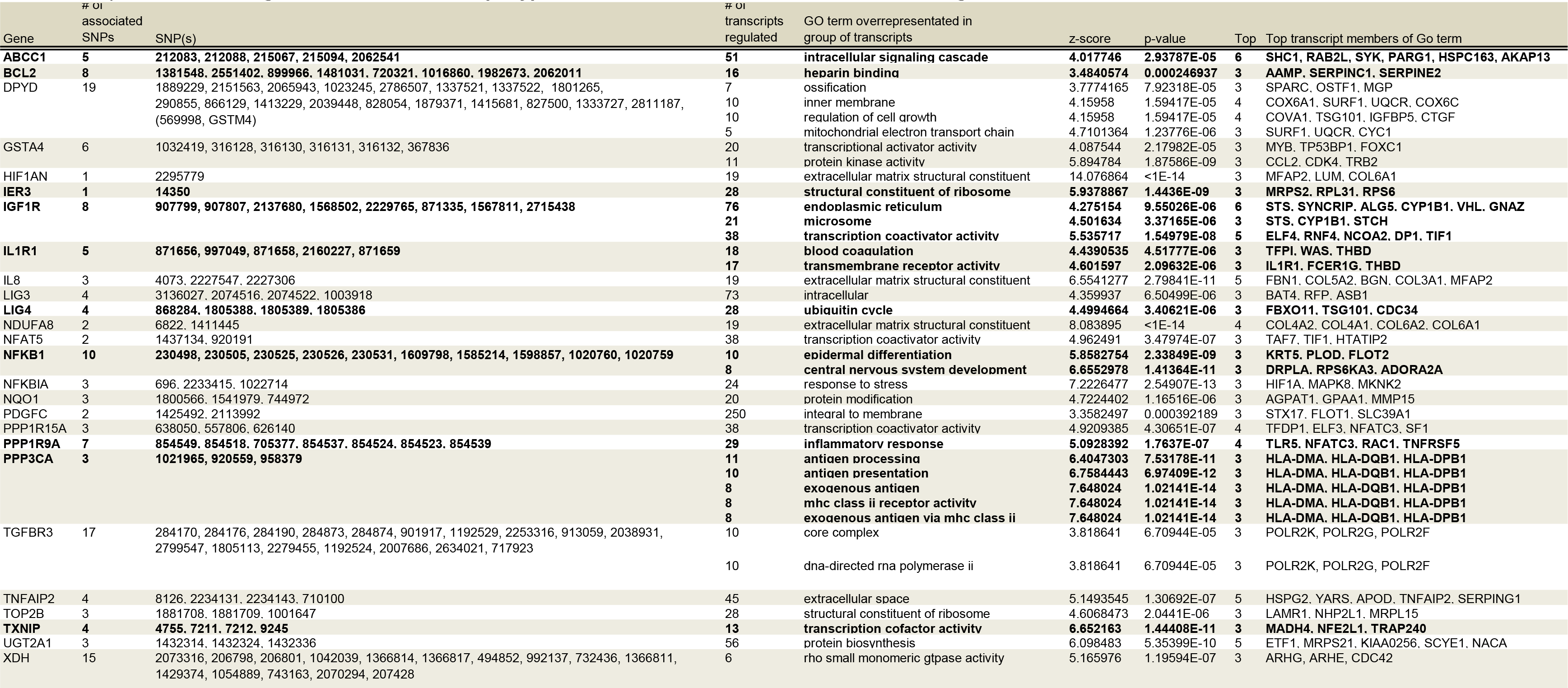
GO analysis of the set of transcripts associated to groups of SNPs in single genes reveals an overrepresentation of GO terms among these transcripts (p-value<0.001, Supplementary Table 5: p-value<0.05). Genes with a significant difference in haplotype distribution between cases and controls are given in bold (Table 3).

In addition, we also found transcripts such as *ANKS1, CREG, NFKB1, TYMS* and *USP1* that were associated each to multiple SNPs **(Supplementary Table 3).**

### Analysis of the haplotype distribution and chromosome wise LD pattern in the case vs. the control population

Haplotypes were estimated for all genes harboring more than 3 SNPs with a maximum distance between neighboring SNPs of 100 kb (n=83). Haplotypes were inferred for the case and control groups separately and the significance of the difference in their distribution was evaluated. A significant difference (p<0.05) in haplotype distribution between cases and controls was observed for 35 genes such as *ABCC1, AKT2, NFKB1, ALOX15B, GSR* and *PIK3CA* **(Table 2).**

**Table 2.**
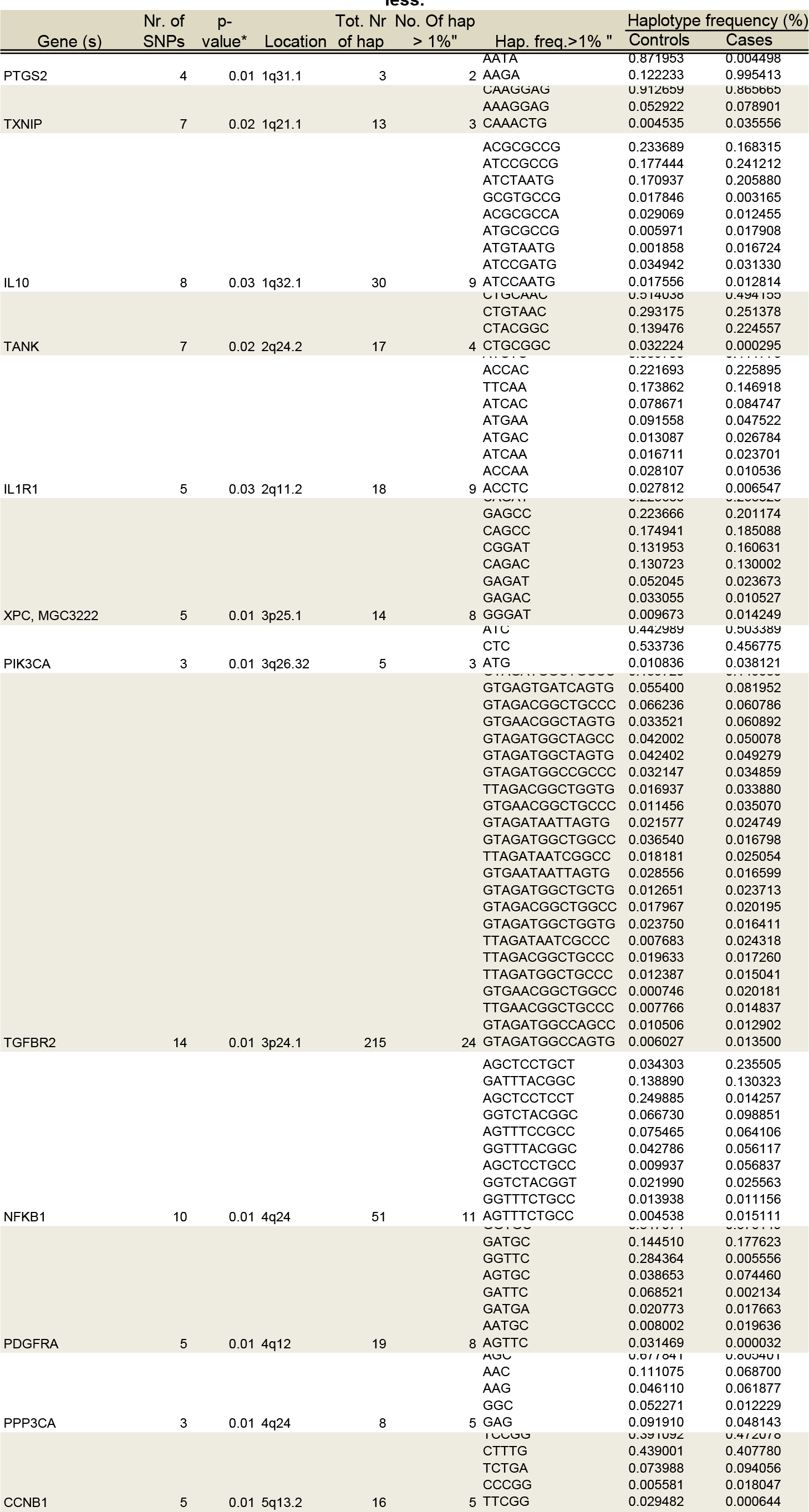
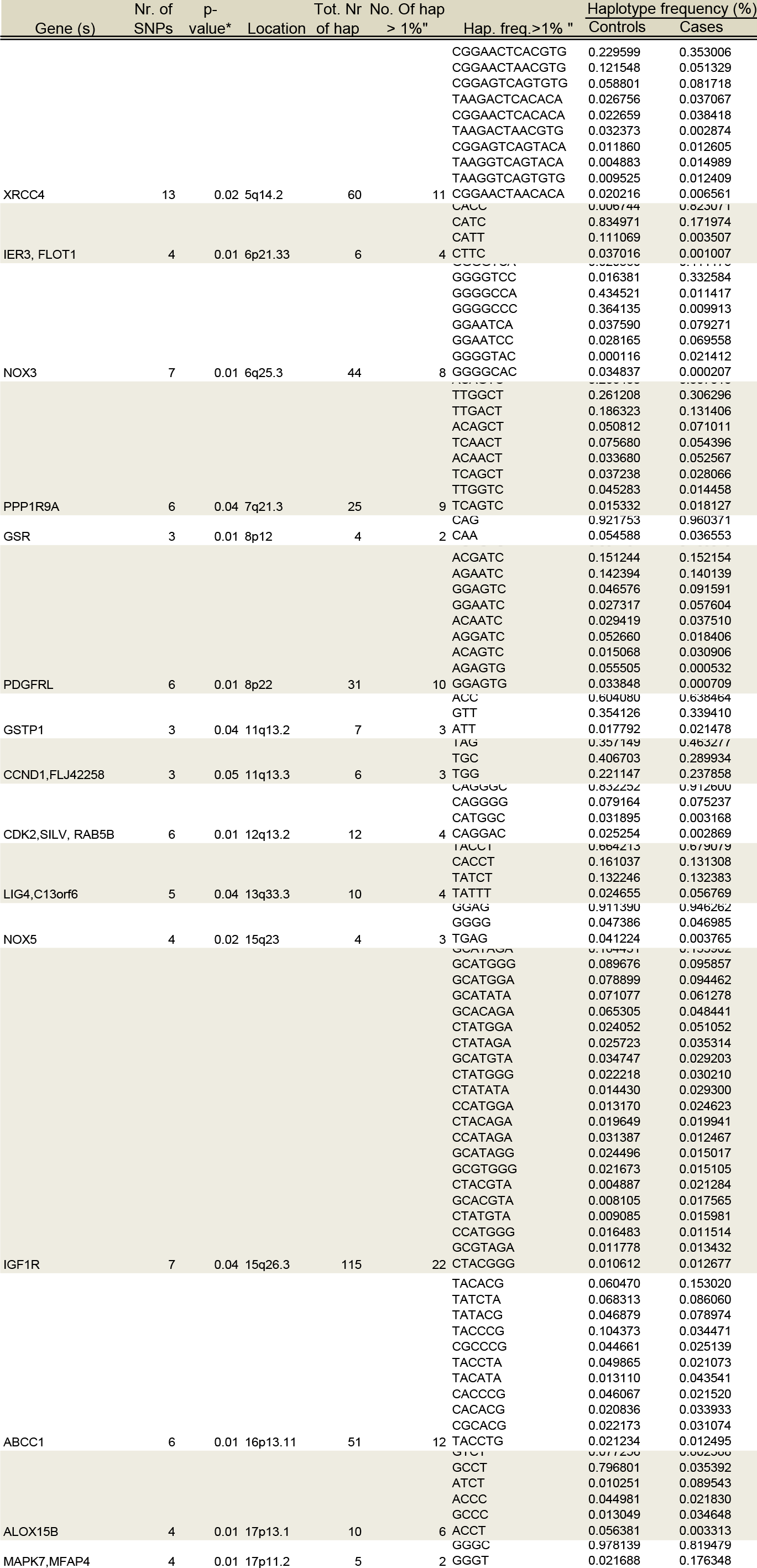
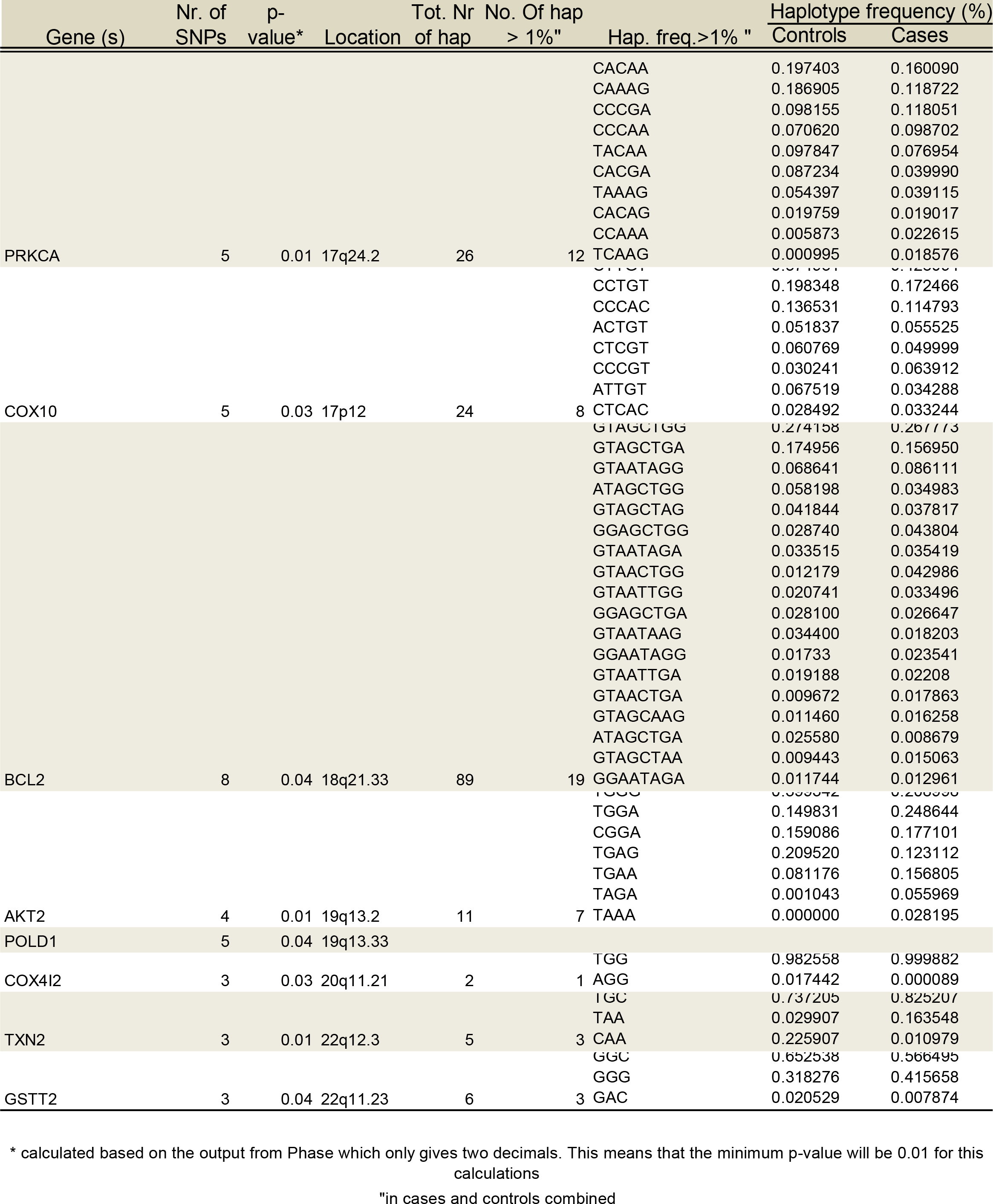
Genes with significantly different haplotype distribution between cases and controls. *P value of 0.01 indicates 0.01 or less.*

The pairwise LD was estimated for: 1) all markers and 2) only between neighboring markers by the standard measurements D’ and r^2^ under consideration of the uncertainty in phase estimation as described in (11). In addition for the neighboring markers, ρ (estimating the population recombination rate across multiple populations) was calculated as described by Evans and Cardon (*9).* Looking at neighboring markers there is a negative correlation (ρ < -0.700) between the LD patterns in cases and controls in 8 genes such as *CDKN1A, EPHX1* and *XRCC1* **(Table 3, panel A).** When including all possible pairwise comparisons for the D’ measure, the Spearman’s correlation analysis revealed a negative correlation for *PQLC2, SOD2* and *PIK3CA* **(Table 3, panel B).** Comparing the pairwise correlation analysis between cancers and controls with the haplotype distribution analysis we see that for the genes where we find a significant different haplotype distribution between cases and controls the correlation is either very low or positive. These results indicate that the difference between cases and controls may be identified by studying together the degree of correlation of LD patterns and the haplotype frequency distribution. Additionally, we investigated neighboring clusters of genes for differences in LD structure and found a negative correlation between the LD values for cases and controls in neighboring regions for gene-pairs such as *IL1A+IL1B, RAF1+XPC* and *NFKBIA+FOS* **(Supplementary Table 4).** These results suggest that the impact of a SNP on susceptibility may be fortified by its organization into haplotype structure including more than one gene, which together may confer higher risk.

**Table 3.**
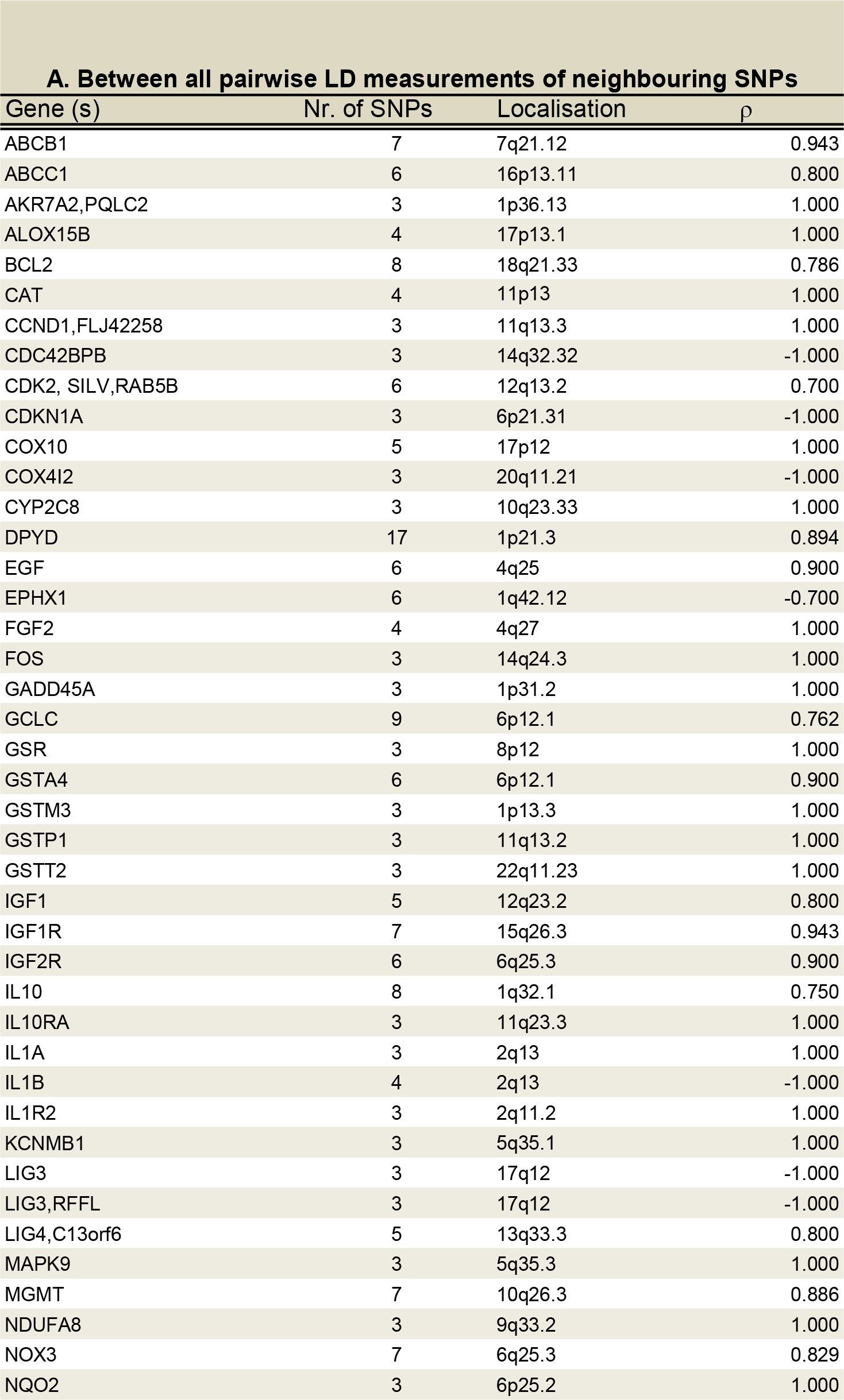
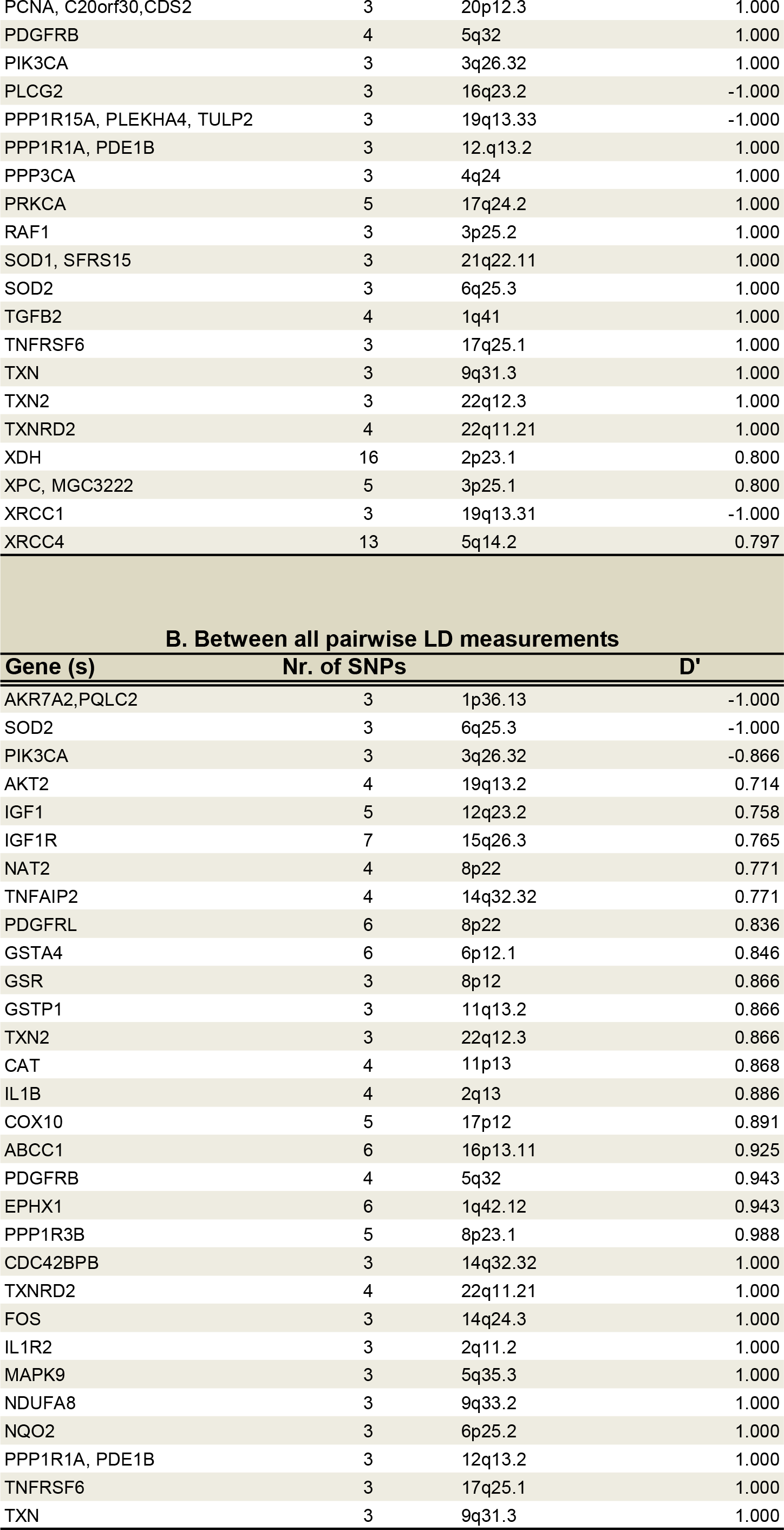
Spearmann’s correlation between LD of cases and controls for neighbouring SNPs (panel A) and all SNPs (panel B) within a gene based on r and D’ values respectively. Listed here are only genes with an absolute correlation between 0.7 and 1

### Impact of the identified putative susceptibility haplotypes on the expression profile;

The haplotypes that were found significantly differently distributed between cases and controls in the genes *ABCC1, BCL2, IGF1R, LIG4, PPP1R9A* and *TXNIP.* were then tested for association to intratumoral expression. The increased complexity with increasing number of estimated haplotypes made it difficult to detect any significant trends for *ABCC1, BCL2, IGF1R* and partly *PPP1R9A* but for both *LIG4* and *TXNIP* a significant association between the expression level of several transcripts and the estimated haplotypes was identified. For TXNIP, the second most frequent haplotype (AAAGGAG, **Table 1)** was found associated to the expression level of *MADH4, NFE2L1* and *TRAP240* (exact p-value <0.001 and 0.001 respectively, **Figure 2a and b**). For *LIG4*, three transcript probes linked to the overrepresented GO term “ubiquitin cycle” were available representing the expression levels of *FBXO11, TSG101* and *CDC34.* Combinations of the second most frequent haplotype (CACCT, **Table 1)** show a significantly different expression level for *FBXO11* (exact p-value 0.009, **Figure 2c).**

**Figure 2.**
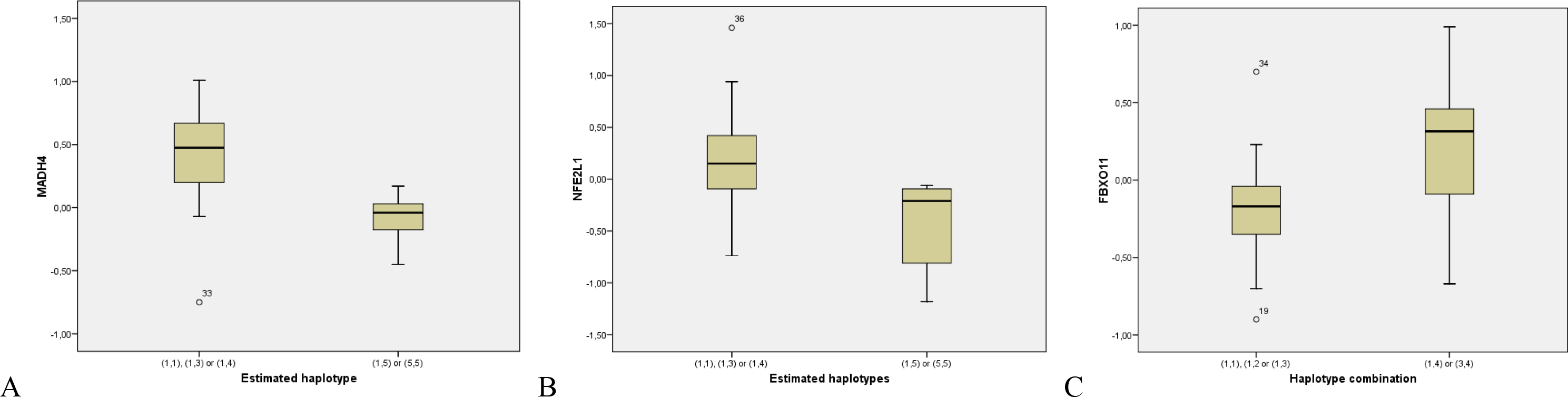
Boxplots showing the spread in the expression levels of the transcripts probes for *MADH4*(A) and *NFE2L1*(B) for the different haplotype combinations of *TXNIP* as well as the spread in the expression levels of the transcripts *FBXO11* for the different haplotype combinations of LIG4

### Frequency distribution of the htSNPs derived from the putative susceptibility haplotypes associated to expression in cases and controls

A total of 42 htSNPs in 9 genes (*ABCC1, IL1R1, PPP3CA, NFKB1, BCL2, IGF1R, LIG4, PPP1R9A* and *TXNIP*) with both significant difference in haplotype distribution between cases and controls and an association between multiple SNPs in the gene an intratumoral expression, either *in cis or trans*, were selected for case control analysis. All in all we genotyped 3484 samples divided in 1592 samples from BC patients/survivors and 1892 controls. 16 of the 42 investigated SNPs were found associated or borderline associated with case-control status **(Table 4).** Three SNPs, rs 215094 in ABCC1,(p<2.25E-04) rs878335 in *IGF1R* (p< 5.58E-09) and rs1805388 in *Lig4 (p<* 7.73E-6) were significant after BonFerroni correction with the SNP in *IGF1R* reaching genome wide significance level.

### Controls vs hapmap Caucasians

Population subdivision between our sample material and the HapMap samples was estimated by the Fixation index (Fst) which measures the population differentiation between two or more (22). The Fst was calculated separately for the nine genes with ≤7 loci available for analysis (*BCL2, IGF1R, IL10, NFKB1, NOX3, TANK, TGFBR2, TXNIP* and *XRCC4*, **Table 1)** and then averaged over all genes. The average F_st_ was 0.0065, indicating a negligible difference between the two populations.

## Conclusion

Several studies have looked into the relationship between single SNPs and risk of sporadic breast cancer both at the single SNP level and the GWAS level. The success of the former in identifying low penetrance alleles have been limited while the latter has identified regions of 10q26 (*FGFR2*), 16q12.1 (*TNRC9*), 5q11.2 (*MAP3KI*), 8q24, 11p15.5, 5q12 and recently 1p11.2, 14q24.1 (RAD51L1), 3p24 and 17q23.2 to be linked to risk of sporadic breast cancer (3,23-27). In this study we have chosen to look at the association between haplotypes and LD patterns in more than 80 genes distributed across all chromosomes and how they differ between cases and controls and identify differences in both, interestingly not at the same time, in important cancer related genes such as *NFKB1, PIK3CA* and *CDKN1A.* We also link the results of our haplotype analysis to our previously published results revealing an association between the germline variation and the expression level in the tumor itself (13). Our SNPs are not representative for the whole genome - they are selected from a candidate gene approach but they anyway make grounds for comparing haplotype patterns between cases and controls and to estimate to what extent these results can be extrapolated to other populations through the genetic similarity with data extracted for the Caucasian samples included in the HapMap project. If we manage to find SNPs in the classical and novel regulatory areas of the genes that correlate to the expression of genes in breast cancer, we will be able to predict the risk of developing certain molecular portraits of breast cancer before the cancer has at all occurred.

## Acknowledgement

This project was supported by The Norwegian Cancer Association (project number D-03067 and D-99061) and The National Programme for Research in Functional Genomics in Norway (FUGE, project numbers 151924/150 and 15204/150), funded by The Research Council of Norway. Genotyping was partly performed at the Uppsala SNP technology platform, which is funded by the Swedish Wallenberg Consortium North (WCN).

## Table legends

Supplementary Table 1. SNPs included in analysis with information on gene affiliation, chromosomal position, allelic variants and strand genotyped.

Supplementary Table 2. Multiple SNPs located in the same gene were found associated to the expression level of a number of transcripts by both ANOVA and QMIS analysis in [1]. Listed here are the gene info, rs-numbers, probe id of associated transcripts, most significant p-value from association analysis as well as whether the association is *in cis* or *in trans.*

Supplementary Table 3 Transcripts associated to genetic variation of multiple SNPs located within the same gene by both ANOVA and QMIS analysis in [1]. Listed here are gene info for identified transcripts,rs-numbers and gene info of associated SNPs, most significant p-value from association analysis as well as whether the association is *in cis* or *in trans.*

Supplementary Table 2. Spearmann’s correlation based on D’ values between LD of cases and controls calculated in the intergenic areas. Listed here are only intergenic regions with an absolute correlation between 0.4 and 1

Supplementary Table 4. List of transcripts regulated by several SNPs

Supplementary Table 5. GO analysis of the set of transcripts associated to groups of SNPs in single genes reveals an overrepresentation of GO terms among these transcripts (p-value < 0.05). “Top” indicates the number of the regulated transcripts associated with the given GO term.

## Figure legends

Supplementary Figure 1 Flow chart of the sample material and analysis

Supplementary Figure 2a-u.Chromosome wise LD for the cases together with information on overrepresented GO terms among associated transcripts. X-axis indicates the significance level of the LD while the |D’| values are plotted on the Y-axis, values along the diagonal are intragenic, adjacent panels give information on intergenic regions.

